# Parrots and Humans Prefer Pitch Sequences of Intermediate Complexity

**DOI:** 10.1101/2025.06.22.660906

**Authors:** Oliver T. Bellmann, Marisa Hoeschele, Melina Witt, Eva Shair Ali, Pui Ching Chu, Roberta Bianco

## Abstract

A well-established inverted U-shape relationship exists between stimulus complexity and human music preference, with preferences peaking at intermediate levels. We investigated whether preference for intermediate complexity is unique to humans or shared with other species. We compared the acoustic preferences of humans and budgerigars, known for their vocal learning and pattern recognition abilities, using a three-option place preference paradigm. Participants were presented with choices between computationally well-defined stimuli varying in complexity: pitch sequences with tones arranged either randomly (high complexity), into repeating patterns with changes (medium), or into a fully repetitive pattern (low). Our results demonstrate that both humans and budgerigars prefer stimuli with intermediate complexity, spontaneously exhibiting an inverted U-shape relationship between complexity and preference without any specific goal or external reward. Because a U-shape relationship has been argued to maximize learning, this preference may be of potential utility for vocal learning, however, further testing could determine whether it is also present for acoustic patterns in vocal non-learning species. Because vertebrates show inverted U-shape responses in a variety of non-auditory contexts, our findings may be rooted in general processes, but may depend on vocal learning to translate to acoustic contexts. This provides exciting opportunities to further investigate the fundamentals of vocal learning within and beyond the auditory and/or musical domain.

**Significance statement:** Appreciating music is part of what it means to be human, but it is unclear why. Comparing budgerigars – a vocal-learning parrot species – and humans, we focused on a sweet spot of musical enjoyment: We like to hear music that’s not too complex and not too simple. Using a place preference setup, we showed that both species prefer to listen to pitch sequences with medium complexity relative to low or high complexity. This shared pattern suggests a fundamental cognitive principle that guides animals to stimuli that maximize learning, promoting vocal communication. Because both budgerigars and humans are vocal learners and music is a socially transmitted behavior, the appreciation of music thus may be rooted in the requirements of vocal learning.

---

A well-established inverted U-shape relationship exists between stimulus complexity (or related factors such as stress or arousal) and preference (“the *U*”, Ten et al., 2024). That is, humans prefer stimuli with medium complexity compared to those with low or high complexity, with preferences changing predictably based on levels of cognitive effort and arousal or interest (Berlyne, 1970, 1971, for a review, see Chmiel & Schubert, 2017).

In non-human animals, the *U* has been observed mainly in learning tasks, but is found also in other behaviors. For example, in ground squirrels (*Spermophilus beldingi*), learning performance follows an inverted U-shaped relationship with cortisol levels: moderate cortisol levels improve the learning of adaptive responses to alarm calls and spatial memory, while learning is impaired at both low and high cortisol levels (Mateo, 2008). Similarly, in rats (*Rattus norvegicus*), stress induced by water temperature affects spatial learning in an inverted U-shape – rats trained at 19°C (moderate stress) made fewer errors in a spatial task compared to those trained at either lower (25°C) or higher (16°C) stress levels (Salehi et al., 2010). Self-grooming behavior in rats was also found to follow an inverted-U shape in relation to anxiety- or stress-like behaviors. In moderately aversive situations (e.g. habituation to a new environment), rats exhibited higher self-grooming behavior compared to very low or highly aversive situations (Fernández-Teruel & Estanislau, 2016). In zebra finches (*Taeniopygia guttata*), the number of heard song exemplars and the quality of song imitation follow an inverted-U relationship (Araguas et al., 2022). Exposure to very few or too many song exemplars yielded poorer imitation scores, while birds exposed to an intermediate number achieved the best imitation, suggesting there is an optimal level of exposure complexity for effective song learning. These findings suggest that the *U* is not limited to humans, but is found in a variety of non-human animals, and corroborate that optimal learning, performance, or behavioral responses are facilitated by moderate levels of stimulus complexity or environmental challenge. Accordingly, the *U* was suggested to be grounded in mechanisms guiding organisms to stimuli maximally useful for learning (Wu et al., 2022; Ten et al., 2024).

Learning also plays a crucial role in many animal acoustic communication systems. Music and bird song, for instance, rely on the repeated social transmission of acoustic patterns (Sierro et al., 2023; Williams, 2006; Rothenberg et al., 2014; Ivanitskii & Marova, 2023). Social learning needed for cultural transmission, here, occurs both horizontally (between peers) and vertically (across generations, Aplin, 2019). Much like birdsong, which is shaped and transmitted through social learning (Williams & Lachlan, 2022), human musical traditions also rely on interaction and imitation (Rehfeld et al., 2021), leading to increasingly complex, diverse, and refined forms over time. However, with each instance of transmission, acoustic patterns also must pass a ‘learner bottleneck’ (Merker et al., 2015). That is, overly complex sequences or configurations are hard for receivers to learn and reproduce, hampering communicative efficiency. Likewise, information that is too simple, such as a pattern highly similar to knowledge already available to or held by an organism, offers little learning opportunity, as research in the visual domain suggests (Wu et al., 2022).

Regarding the acoustic domain, the *U* indeed plays an important role in music which is transmitted via social learning and often relies on repetitive patterns (Deutsch et al., 2011; Fitch, 2006; Margulis, 2013, 2014, 2018; Margulis & Simchy-Gross, 2016; Nettl, 1983; Ockelford, 2017; Simchy-Gross & Margulis, 2018; Tierney et al., 2018). For example, research using iterated learning paradigms shows that initially random sequences of note onsets (i.e., rhythm; Ravignani et al., 2016; Jacoby et al., 2021; Jacoby & McDermott, 2017) or pitches (i.e., melodies; Verhoef, 2014; Verhoef & Ravignani, 2021; Anglada-Tort et al., 2023) gradually condense into sets of discrete and small-integer-ratio elements and get less random. This process, improving signal quality and ultimately facilitating learnability, highlights how music is shaped by both cultural evolution and cognitive biases. Indeed, humans prefer music with a balance of predictability and novelty, avoiding overly simple and overly complex stimuli during both perception and cultural transmission (Delplanque et al., 2019; North & Hargreaves, 1995; Burke & Gridley, 1990; Cameron et al., 2023; Matthews et al., 2019; Witek et al., 2023; Cheung et al., 2019; Bianco et al., 2019; Gold et al., 2019; for a review, see Chmiel & Schubert, 2017).

This *U* might be the result of recent cultural phenomena or a human adaptation specific to music. However, it could also be an adaptation facilitating the cultural transmission of acoustic patterns based on social learning more broadly, which is crucial to music and other non-human communication systems, and thus might also be broadly shared across vocal learners (i.e., species that can modify their vocalizations based on auditory experience), such as birds.

Because the U is discussed as a general learning phenomenon that can be found in many species and is also found in human music, we hypothesize that, in vocal learners, medium complexity auditory signals – being sufficiently structured – might be a good compromise to facilitate yet not oversimplify communication and optimize learning. These signals might therefore be preferred over signals of low or high complexity. It is, however, unknown whether birds and other animals exhibit the *U* for acoustic patterns.

Here, we thus investigated if preference for intermediate complexity in acoustic patterns is unique to humans or if it is shared with another vocal learning species: the budgerigar (*Melopsittacus undulatus*). Budgerigars are parrots known for their vocal learning abilities and pronounced pattern recognition skills (Cai et al., 2018; Fishbein, 2022; Spierings & ten Cate, 2016; Stenette et al., 2023; Tu & Dooling, 2012). They have been shown to prefer rhythmically regular acoustic patterns over random ones (Hoeschele & Bowling, 2016) and vocalize using socially transmitted pitch sequences (Brittan-Powell et al., 1997; Farabaugh et al., 1994).

If the inverted U-shaped preference to acoustic stimuli arises from mechanisms and selective pressures shared by vocal learning species or shared more broadly in the animal kingdom, budgerigars should exhibit a preference for intermediate levels of complexity in pitch sequences. Conversely, if this preference is an adaptation specific to music processing and/or the cultural transmission of music, it should be present in humans but not in budgerigars.

We measured the preferences of budgerigars and humans to pitch sequences using a three-option place preference paradigm. Both species were presented with the same stimuli to ensure maximum comparability. In a given session, listeners were presented with choices between two of the three types of stimuli: Either tones arranged randomly, and/or tones arranged into tone patterns that changed every four repetitions, and/or a repetitive tone pattern that always stayed the same. The third option was always silence as the control condition.

## Methods

### Stimuli

Stimuli were sequences made from permutations of a set of 10 rms-normalized 22-ms harmonic tone-pips. Each pip was made of three simultaneous sine tones that represented the 4th-6th harmonic of a given fundamental frequency. Ten fundamental frequencies were used that were equally spaced on a logarithmic scale between 157.5 Hz and 1250 Hz (12 % steps), such that sine 1 = 4f0 | sine 2 = 5f0 | sine 3 = 6f0 (e.g., 630 | 787.5 | 945 for f0 = 157.5; Supplementary Table 1). At 70 dB SPL, these sounds lie well within the human and budgerigar hearing range (Dooling & Saunders, 1975; Farabaugh et al., 1998; Hefner & Hefner, 2007). All sounds had 3-ms onset and offset amplitude ramps to avoid transients (Okanoya, 2000; Hoeschele et al., 2013).

Sequences of the 10 pips were then created such that there was 3 ms of silence between each pip. This was done using three types of permutation procedures corresponding to the three conditions *Random, Regular*, and *Regular with changes*. Every sequence consisted of 800 sounds for a total of 20s, and each of the 10 possible pips occurred 80 times. We created 5 different 20-second sequences for each condition.

For the *Random* (Ran) sequences, the 10 pips were concatenated in random order, then all 10 pips were again concatenated in a new random order, etc. Each tone-pip thus occurred with equal probability across the entire sequence.

The *Regular* (Reg) sequences consisted of the 10 pips from the set, concatenated in a random order, and then this order was repeated for 80 identical cycles.

For *Regular with changes* (VarReg), a pattern of 10 pips was repeated four times, then, a different pattern was repeated four times, and so on, until 20 different repeated patterns had occurred.

### Information-theoretic modelling

To obtain a quantitative information-theoretic measure of complexity, stimuli were characterized using the Prediction by Partial Matching-decay (PPM-decay) model (Pearce, 2018, Harrison et al., 2020), which expresses tone-by-tone unexpectedness in terms of Shannon’s information surprise – or information content, IC – conditioned upon the preceding musical context. PPM-decay was implemented in R (Harrison et al., 2020, https://github.com/pmcharrison/ppm) based on a variable-order Markov model framework that learns the stimulus statistics by analyzing symbolic sequences event by event. As it processes the sequence, the model stores the occurrences of events and subsequences of events (*ngrams*), and it generates a conditional probability distribution for each tone given the preceding tones. Thus, for each event in a sequence, the model quantifies its IC using the probability distribution of previously encountered observations. Contrary to previous learning approaches with perfect memory, PPM-decay also features a number of memory parameters that determine the weighting of historic data, improving its biological plausibility. Here we used its short-term memory configuration, whereby the model learns only from the current 800-sound sequence. Averaging the IC across all events in a sequence provides an index of the sequence’s overall predictability. We interpret this as a measure of complexity, predicting that VarReg sequences (with medium information content) will be perceived as more complex, hence more preferable, than Reg ones (with low information content) and less complex than Ran ones (with high information content).

As shown in Figure 1, the model was run on the full stimulus set. A Shapiro-Wilk test suggested the data were not normally distributed (p < 0.001), therefore, a Friedman test was conducted, indicating significant differences between the conditions (p = 0.007). Subsequent Wilcoxon tests showed that IC was significantly higher in Ran than VarReg and Reg (mean IC = 3.5, 1.867, and 0.346, respectively; all Wilcoxon, p = 0.043), confirming that conditions presented three clearly separable levels of complexity: high, medium, and low.

**Figure 1.**
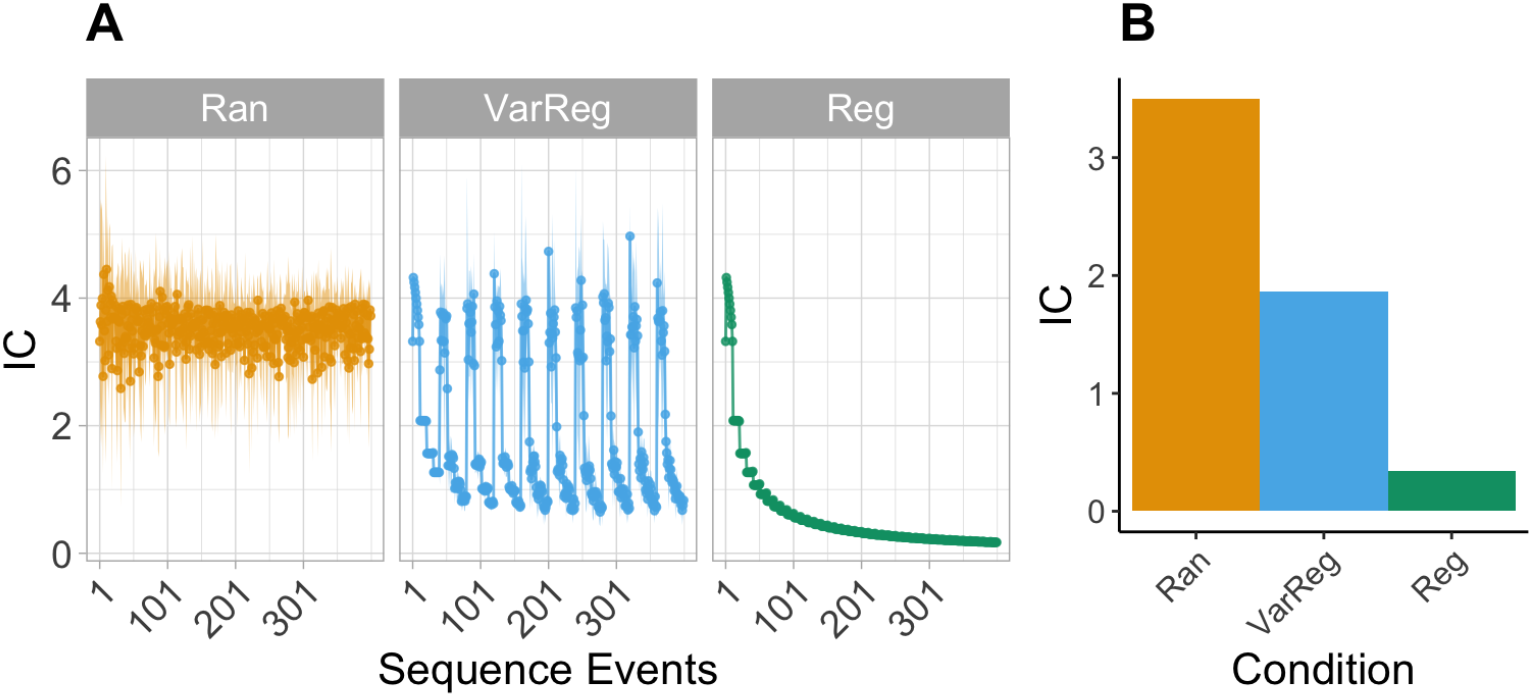
Information content (IC) of the First 400 Events (A) and mean Information Content by Condition (B) across the Ran, Reg, and VarReg Sequences.

### Participants

We tested n = 12 adult budgerigars (6 female, 6 male) housed in the budgerigar lab and n = 32 normal-hearing adult humans (16 female, 16 male; mean age = 27.66 years (20-43), SD = 4.81 years). Both species were tested at the Acoustics Research Institute of the Austrian Academy of Sciences in Vienna, Austria. Human participants were compensated for participation in the experiments in the form of 10 euros (€) per hour of participation.

### Apparatus

We individually assessed the time participants spent with the sequence types as a proxy of preference using a place preference setup (see Wagner et al., 2020).

For budgerigars, this setup consisted of an aviary with three perches as shown in Figure 2A, each fitted with an infrared barrier and adjacent speakers (Vifa full-range speaker 10 BGS 119/8; 104 mm full-range loudspeaker, 80-16000 Hz frequency range, 86 dB characteristic sound pressure), which were driven by amplifiers adapted from Edirol MA-5D active loudspeaker systems (Majdak et al., 2010). Sitting on a perch disrupted the infrared barrier, and playback of sound from the adjacent speaker was triggered after > 300 ms disruption (avoiding triggers by fly-throughs). Stimuli were presented at 70 dB SPL at the middle of the perches (digital sound level meter: AZ Instrument, AZ 8922). During every session, a fourth speaker (M-Audio AV40) with an MP3 player (HIFI WALKER MP-7 8 GB) under the testing aviary played a recording from the bird’s home aviary at 50 dB to alleviate stress from isolation during testing. Water and food were available at the bottom of the testing aviary ad libitum during sessions.

**Figure 2.**
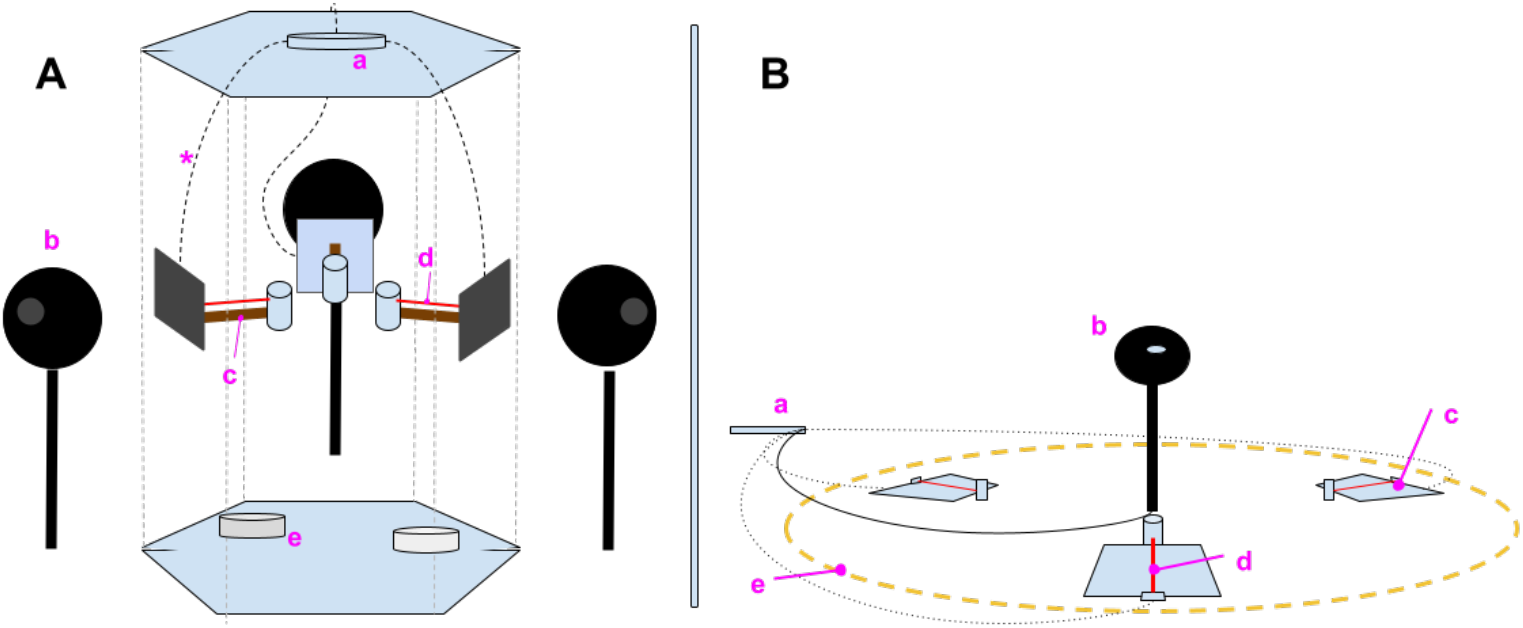
Place Preference Apparatuses for budgerigars (A) and humans (B). Note. **A:** Hexagonal aviary for testing place preference in budgerigars. **B:** Human place preference arena. a = arduino®; b = speakers; c = perch (**A**), place preference mats (**B**); d = infrared beam; e = food and water bowls (**A**), circular marking on the floor (**B**).

For humans, the setup consisted of three floor mats with shoe icons, in a yellow circle (Figure 2 B). In the circle’s center was a speaker (Vifa full-range speaker 10 BGS 119/8; 104 mm full-range loudspeaker, 80-16000 Hz frequency range), whose playback level was set to 55 dB SPL at the center of the mats. Standing on a mat disrupted an infrared beam, which triggered sound playback from the central speaker if the participant stayed for at least 300 ms.

### Procedure

For birds, sessions were carried out in the hexagonal aviary (Figure 2A; see also Wagner et al., 2020). For each 2-hour session, one speaker played one sequence type (e.g., Ran), another played a different sequence type (e.g., VarReg), and the third remained silent. If a listener stayed on a perch (which triggered sound playback) beyond the duration of a stimulus WAV file, a new sequence from the same condition was played. The software cycled through all stimuli from the respective condition in a randomized order until all were played, then repeated the cycle randomly. Leaving the perch would fade out the sound over a 5-ms period, returning to a perch for at least 300 ms triggered the playback of a new stimulus. The program recorded perch condition, time spent on the perch, stimulus playback duration, and the number of > 300-ms perch visits. Following each session, birds were returned to their home aviary. For each bird, multiple sessions were conducted across different days, such that all of the sequence/perch combinations (e.g., Ran/Reg/Silent) were assessed three times for a total of nine sessions. Across sessions and birds, the positions of sequence types were counterbalanced to minimize effects from perch location biases (Supplementary Table 2).

For humans, the procedure was similar, except that first, the participants were told that their task was to explore the area inside a yellow circle for 5 minutes, that there would be 9 such 5-minute sessions, that they could take breaks ad libitum, and that they could end their participation at any point. Next, participants read and signed a consent form prior to testing. After the sessions, participants completed a listener survey, which included demographic questions. Then, they also listened to the sequences heard during the place preference experiment on headphones (Sennheiser HD 280 pro, 64 Ω) and rated them on predictability, interest, and liking, using 7-point visual analogue scales (1 = “Not at all”, 7 = “Very”). Any remaining questions revealing the purpose of the experiments were answered after completion of the surveys.

### Data Analysis

We analyzed the data using R (4.4.1, 2024-06-14) and RStudio (2024.04.2+764, 2024-06-10), employing linear mixed-effects models (lme4 package, Baayen et al., 2008) with REML estimation. All models included subjects as a random effect. The fixed effects included condition (Ran/Reg/VarReg/Silence) and sex (Male/Female), as well as their interaction. Session order was included as a covariate of no interest. Statistical significance was assessed through likelihood-ratio tests conducted using the ‘anova’ function in the stats package. Follow-up contrasts were conducted using the ‘emmeans’ package and the Tukey method to account for the increased risk of type I error resulting from multiple comparisons. The adjusted p-values were calculated to determine significant differences between conditions, with significance level of α = 0.05.

## Results

As shown in Figure 3, our analysis investigated how the amount of time spent on the perch per trial differed between the four conditions (Ran, VarReg, Reg, Silence). A trial is defined as every instance in which a perch (for birds) or a mat (for humans) was contacted such that the infrared barrier was disrupted for longer than 300 ms. For the relative mean average time the individual budgerigars spent with the different conditions, see Supplementary Figure 1.

**Figure 3.**
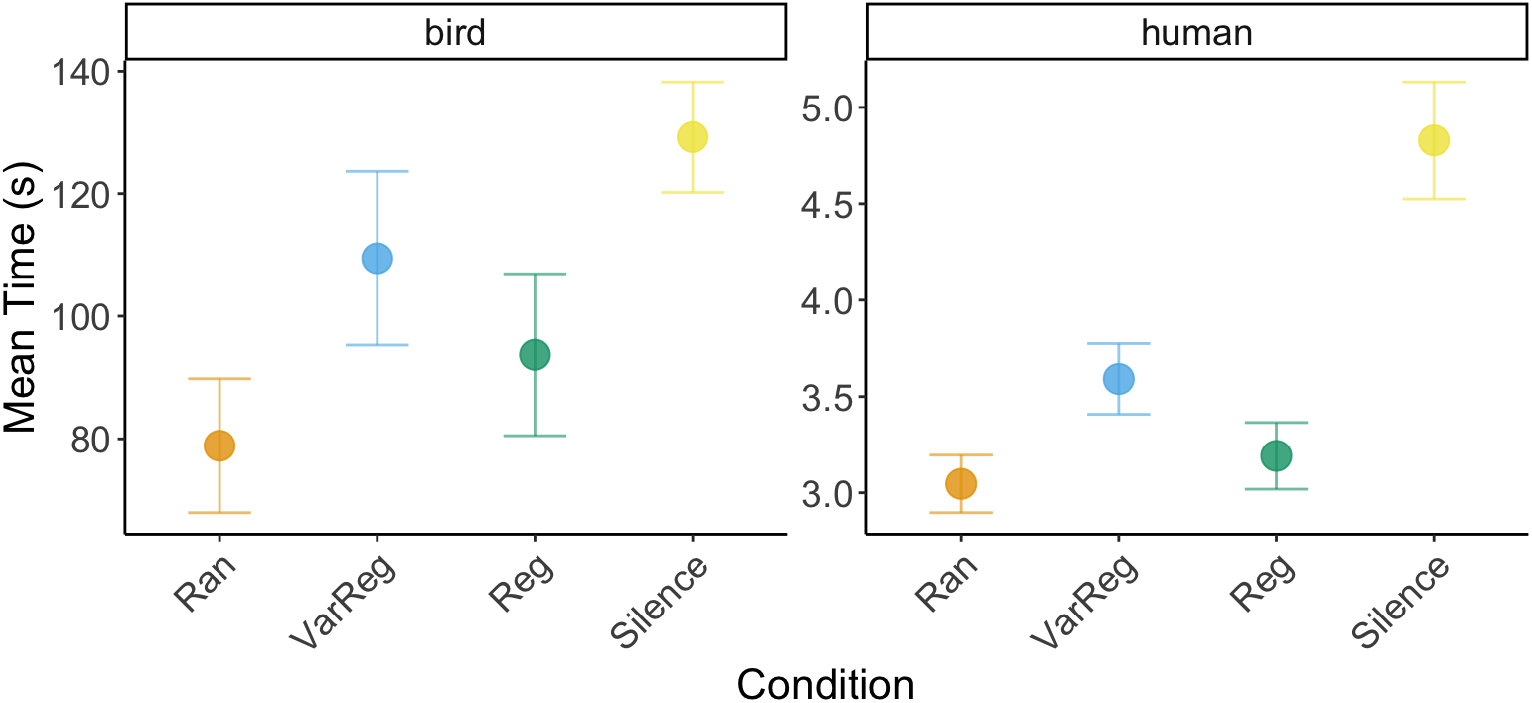
Mean time (in seconds) and standard error of the mean across subjects and trials for the four conditions in budgerigars (A, left) and humans (B, right), Note. Y-axis: Mean time per trial in seconds. X-axis: Ran = random pattern; VarReg = partially repetitive pattern; change of the pattern every 4 repetitions; Reg = single repetitive tone pattern. For a given session, only three of the four total conditions were available (e.g., Ran, Reg, and Silence) and Silence was always present as a control condition.

For birds, the model revealed significant main effects of condition (χ^2^(3) = 15.11, p = 0.002) and session (χ^2^(1) = 9.15, p = 0.002), but not sex (χ^2^(1) = 0.003, p = 0.957). The interaction included in the model – between condition and sex – was not significant (χ^2^(3) = 5.53, p = 0.137). This suggests that overall time spent differed across conditions, with this effect not varying by sex.

A post-hoc analysis with pairwise comparisons indicated no statistically significant differences between Ran, VarReg, and Reg (all ps > 0.391). However, whilst birds spent more time on Silence than both Ran (p = 0.003) and Reg (p = 0.042), we observed no significant difference between Silence and VarReg (p = 0.374). This suggests that the budgerigars spent significantly more time with Silence compared to Ran and Reg, but similar time between Silence and VarReg.

For humans, the LMM revealed no significant main effects of condition (χ^2^(3) = 1.601, p = 0.659) or sex (χ^2^(1) = 0.924, p = 0.336). However, there was a significant interaction effect between condition and sex (χ^2^(3) = 13.30, p = 0.004), suggesting that the effect of condition on time spent differed for males and females.

We thus carried out separate analyses of males and females using a LMM as described above (See Supplementary Figure 2). There was a significant main effect of the condition for males (χ^2^(3) = 17.1759, p < 0.001), but not for females (χ^2^(3) = 1.7592, p = 0.6238).

A post-hoc analysis of the male data with pairwise comparisons showed that males spent significantly more time with VarReg than with all other conditions (VarReg vs Reg, p = 0.035; VarReg vs Ran, p = 0.024; VarReg vs Silence, p < 0.001). The other contrasts yielded no significant differences (Ran vs Reg; Ran vs Silence; Reg vs Silence; all p > 0.327).

### Listener Survey Results

Figure 4 shows the listener survey results.

**Figure 4.**
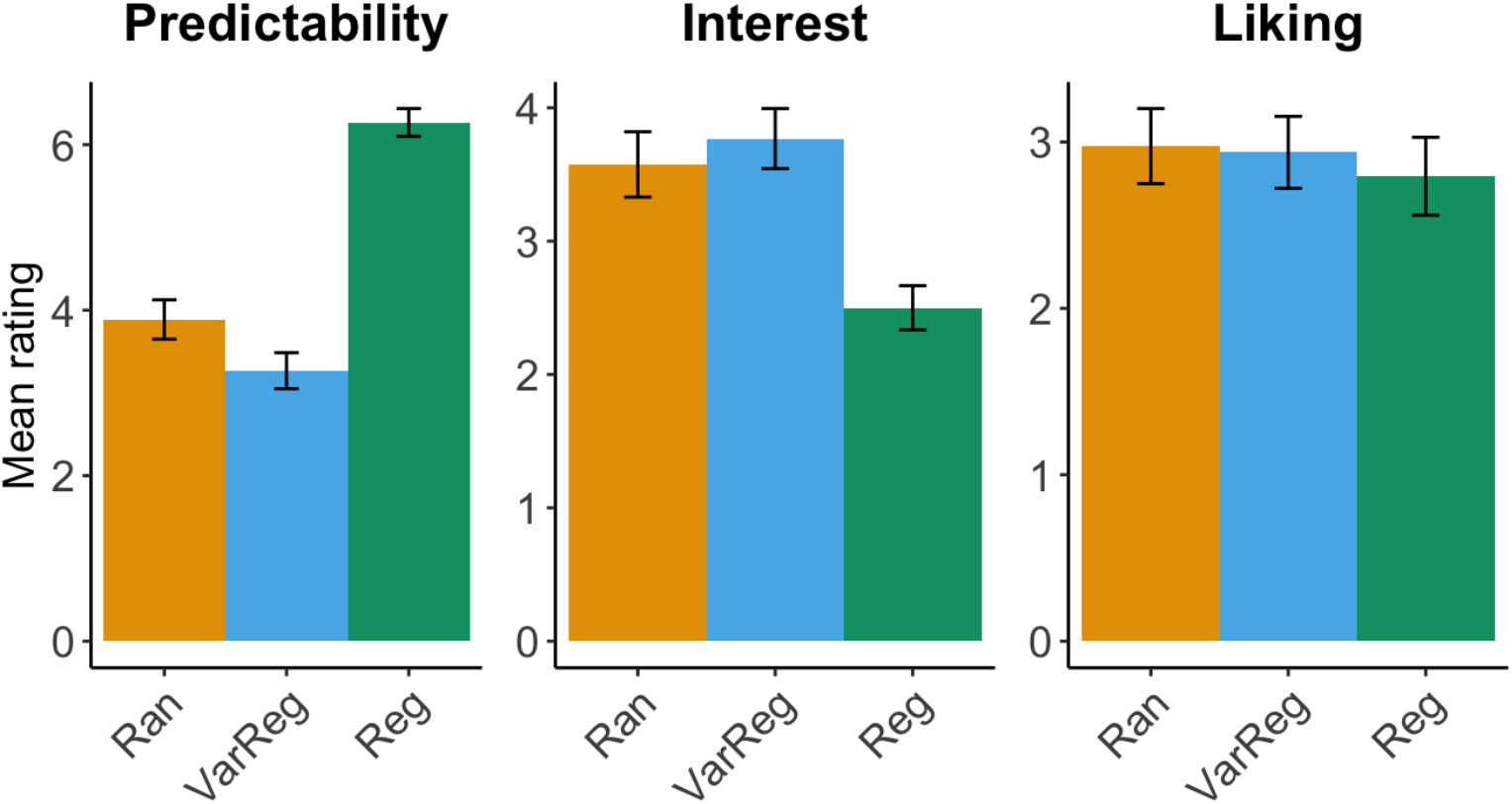
Listener survey results: Mean rating per condition (x axes) on a scale of 1-7 and standard error of the mean across subjects.

Concerning predictability, linear mixed-effects models showed significant main effects of the sequence type (χ^2^(2) = 253.324, p < 0.001), no effect of sex (p = 0.3477), and no interaction (p = 0.0824). A post-hoc analysis with pairwise contrasts revealed that predictability was greater for Reg than VarReg (p < 0.0001) and Ran (p < 0.0001), and greater for Ran than VarReg (p = 0.0001).

In the model for interest, there was a main effect of the sequence type (χ^2^(2) = 23.563, p < 0.001) and sex (χ^2^(2) = 6.849, p = 0.009), but not of their interaction (p = 0.797). Pairwise contrasts revealed that compared to Reg interest ratings were greater for VarReg (p < 0.0001), as well as for Ran (p < 0.0001), while there was no difference between VarReg and Ran (p = 0.5532).

In the model for liking, there was no effect of sequence type (χ^2^(2) = 1.0083, p = 0.604), no effect of sex (p = 0.0533), and no interaction (p = 0.9399).

Next, we investigated the relationship between ratings and IC using linear mixed models. The IC significantly predicted ratings of both predictability (χ^2^(1) = 177.72, p < 0.0001) and interest (χ^2^(1) = 62.538, p < 0.0001), but not liking (p = 0.1246). To assess whether models that allow for a curved relationship better predicted the ratings, we introduced quadratic terms for IC to our models, which also showed a significant main effect of IC on ratings of predictability (χ^2^(2) = 485.32, p < 0.0001) and interest (χ^2^(2) = 114.39, p < 0.0001), but not liking (χ^2^ (2) = 2.6535, p = 0.2653). Comparing the linear and quadratic models using ANOVA with maximum likelihood instead of REML showed that the quadratic models were a significantly better fit for both predictability (p < 0.0001) and interest ratings (p < 0.0001). This suggests that predictability and interest ratings may change with IC up to a certain point, and then potentially reverse or plateau, as shown in Figure 5.

**Figure 5.**
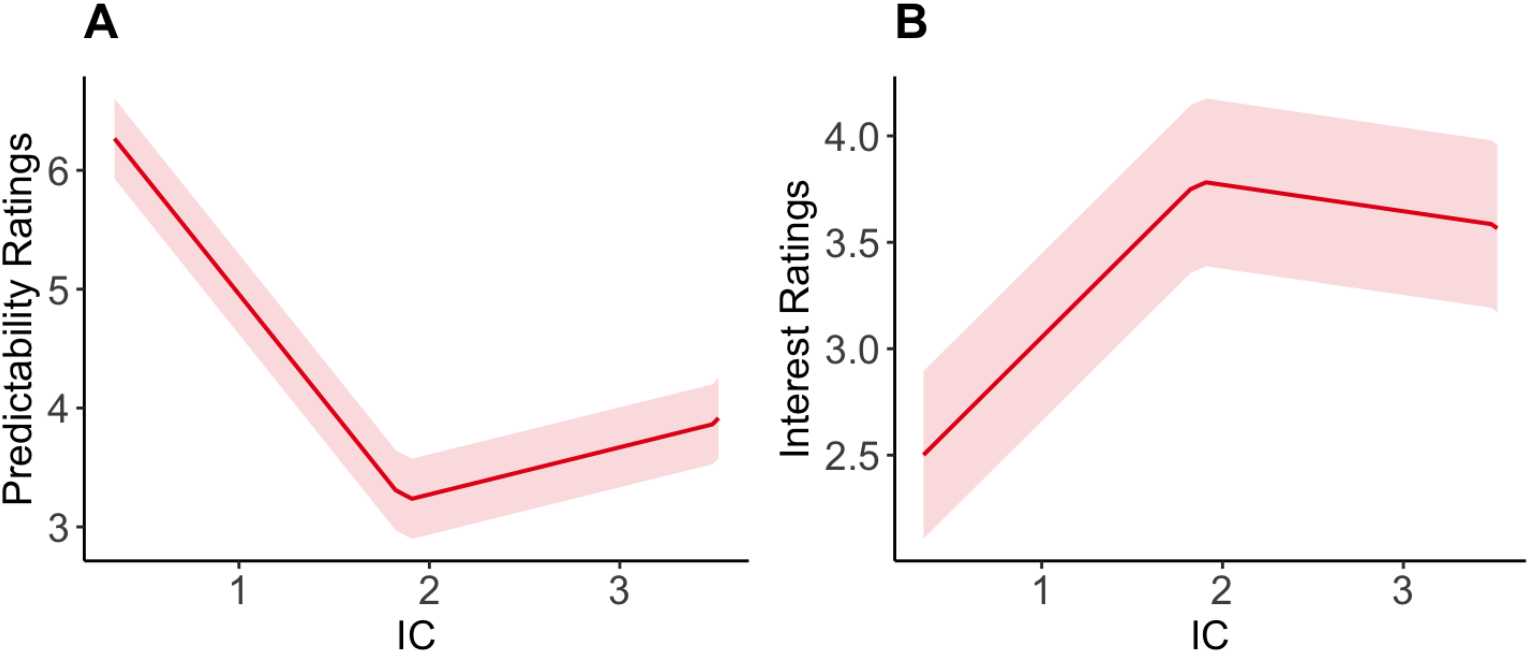
Predicted ratings of predictability (A) and interest (B) with confidence intervals.

## Discussion

We investigated the relationship between stimulus complexity and preference in the auditory domain across two vocal learning species: budgerigars and humans. This relationship was generally proposed to be shaped by mechanisms that preferably draw organisms to intermediate complex information optimal for learning. We investigated whether such an inverted-U shape relationship (The *U*) applies to the auditory domain in vocal learners, because their communication greatly depends on efficient learning in the auditory domain. Using pitch sequences of low, intermediate, and high complexity, we tested preferences in budgerigars and humans, two vocal learning species, each with communication systems that rely on socially transmitted pitch sequences.

Our principal finding is that both budgerigar and human preference for pitch sequences exhibit the *U*, as shown by behavioral responses in both species and by survey responses in humans. This presents a novel synthesis of the relationship between human music and animal cognition in terms of the U. The similarity of our results across two genetically distant species of vocal learners could reflect behavior guided by a similar optimized learning strategy for music and bird communication, two widely recognized model systems for vocal learning (e.g., Patel, 2021; Rothenberg et al., 2014). This strategy may guide learning by channeling cognitive resources towards auditory material based on its statistical properties. Similar to the visual domain (Wu et al., 2022), it may prioritize material of medium complexity that balances novelty and familiarity: complex enough to offer new information and build new models of the world, but also sufficiently related to already existing models to enable their consolidation and refinement. Learners have to deploy their limited resources optimally, because it is impossible to use all available information. Likewise, ignoring opportunities for learning hampers the improvement and expansion of existing skills and knowledge (Wilson et al., 2019; Wu et al., 2022). The suggestion that there are mechanisms guiding learners to stimuli useful for learning is supported by evidence from an investigation of in-ovo sound processing in birds. Compared to vocal non-learning species, vocal learners show stronger physiological responses to conspecific vocalizations (i.e., increases in heart rate) than vocal non-learning species already in ovo (Colombelli-Négrel et al., 2021). This heightened response strength suggests that perhaps vocal learners, compared to non-learners, have a broader and/or additional set of mechanisms that first guide attention to important stimulus types before filtering them according to complexity.

However, because we did not test vocal non-learners, it is still unknown whether the U is also present in these species. In general, vertebrates show an inverted U-shape relationship in a variety of behavioral, non-auditory contexts, for example, between prey size and preference of prey choice (Metcalfe & Jacobs, 2010), the familiarity of playmates (Ham & Pellis, 2023), or the level of surprise in sequences of visual shapes (Wu et al., 2022). Concerning the acoustic domain, research in mice showed that moderate states of arousal are associated with better performance for both categorizing polyphonic melodies in a discrimination task (Hulsey et al., 2024), and for a tone-in-noise detection task (McGinley et al., 2015). This data suggests that we would find similar results in vocal non-learners. Further testing on whether a U-based learning strategy in the auditory domain is shared beyond vocal learners and also investigating the attraction to sounds in relation to stimulus complexity in more species could be fruitful avenues for research into auditory cognition across the animal kingdom.

The overall attraction of birds and humans to patterns that are likely better for learning indicates links of the U to aesthetic phenomena. In general, stimuli that are more appealing may drive animal behavior for a variety of functional goals, often linked to social learning, such as mate choice (MacLaren, 2024) and nest material selection (Guillette et al., 2016). The U we observed here may be a driving force underlying aesthetic cognition. Aesthetic cognition, the capacity to perceive and create beauty, as well as making choices based on aesthetic judgments, is potentially shared across many species (French, 2022; Welsch, 2004; Westphal-Fitch & Fitch, 2018). Our experiment, being designed for cross-species use, could be used for large-scale cross-species investigations, pushing that line of inquiry by further examining the relationships between complex auditory stimuli and preference.

A few things about our human results are also worth noting. For example, our additional sound rating task with humans revealed that high complexity random sequences were rated as more predictable than our medium complexity sequences that repeated and then changed, despite the higher IC of high complexity (Ran) sequences. High complexity sequences were potentially characterized by being predictably unpredictable, a phenomenon that can lead to low uncertainty despite fluctuating information (Shin & DuBrow, 2021). To illustrate this with an example, the individual sounds of the drops that constitute the event type “Rain” are impossible to time, yet the sound of rain itself is likely not perceived as unpredictable, but rather soothing and constant. In contrast, with the medium complexity (VarReg) sequences, only the pattern change itself was predictable, occurring after every four repetitions. The contents of the patterns (i.e., both their contour and the order of individual tone-pips), however, were randomized anew with every change of the pattern, possibly leading to a perceived uncertainty higher than that of the high complexity sequences.

Notably, interest and predictability ratings mirrored each other. In other words, medium complex stimuli were rated the most interesting and the least predictable. Conversely, low complexity stimuli were rated the least interesting, while high complexity stimuli obtained intermediate ratings in both interest and predictability.

Previous research has highlighted that the relationship between aesthetic judgements and predictability is complex and dependent on the immediate context auditory stimuli are situated in (Albury et al., 2023). Rather than preferring information that is predictable, humans prefer information that is predictive of other events in the immediate context (Braem & Trapp, 2019). This suggests that people find stimuli that are predictive of other events more interesting or attention-worthy than stimuli that are predictable but are not predictive of other events. In our stimuli, the internal structure of the medium complexity sequences and their predictiveness might explain the higher interest ratings for these sequences. Four repetitions of a randomly assembled tone sequence would trigger the start of a new random sequence to be repeated 4 times, and so on. The medium complexity stimuli thus can be regarded as a succession of events, that, like a countdown, indicated the impending change of event to a new sequence, which again clocked down four times to again trigger the event of a change in pattern to a new sequence. Therefore, the medium complexity sequences were rather unpredictable in terms of their content, yet their subordinate events were highly predictive of other events, that is, the change of pattern.

Taken together, preferences for predictive information may explain the complementarity of the U-shaped predictability ratings and the inverted U-shaped interest ratings, while also accounting for the human and budgerigar results from the place preference apparatus. Because preference of predictive stimuli is found in non-human primates (Smith & Beran, 2020; Smith et al., 2017), pigeons (Roper & Zentall, 1999), and rats (Cunningham & Shahan, 2019), our results suggest that investigating the propensity for predictive stimuli, in conjunction with preference for intermediately complex stimuli across a wider range of animals might be an exciting avenue for research into the foundations of aesthetic perception across the animal kingdom.

## Ethics Statement

The place preference paradigm used in this study is a noninvasive experimental technique and was carried out according to Austrian animal protection and housing laws, and was approved by the ethical board of the behavioral research group in the faculty of Life Sciences at the University of Vienna (approval number 2015-005). The participation of individual birds in the experiments was approved by an animal keeper at the Acoustics Research Institute. All procedures involving humans were approved by the Commission for Science Ethics of the Austrian Academy of Sciences (Statement of the Commission for Science Ethics of the Austrian Academy of Sciences of April 23, 2024). In the experiments involving humans, we followed the European Charter of Fundamental Rights, worked along the guidelines of “Good Scientific Practice,” and fulfilled the ethical principles for research involving human subjects (Helsinki Declaration).

## Acknowledgements

We would like to thank Barbara Götz and Barbara Timmler for taking care of the animals, Moritz Müller for helping with the human experiments, and we would like to thank our human and non-human participants.

## Data Availability Statement

The datasets for this study can be found in the OSF repository (link: https://osf.io/scdwm/?view_only=6212f56b481249eb973e945214f8a948)

## CRediT Author Statement

**Oliver Tab Bellmann**: Investigation, Data Curation, Resources, Formal Analysis, Visualization, Writing - Original Draft, Writing - Review & Editing, Project administration

**Marisa Hoeschele**: Conceptualization, Supervision, Writing - Review & Editing, Methodology

**Melina Witt**: Investigation, Data Curation, Validation, Writing - Review & Editing

**Eva Shair Ali**: Investigation, Writing - Review & Editing

**Pui Ching Chu:** Investigation, Data Curation, Writing - Review & Editing

**Roberta Bianco**: Conceptualization, Supervision, Writing - Review & Editing, Software, Methodology, Formal Analysis, Visualization, Validation

## Supplementary materials

**Supplementary Table 1.** Sine triads used in the pitch sequences.

| Triad no. | f sine 1 (Hz) | f sine 2 (Hz) | f sine 3 (Hz) | virtual f0 |
| --- | --- | --- | --- | --- |
| 1 | 630 | 787.5 | 945 | 157.5 |
| 2 | 794 | 992.5 | 1191 | 198.5 |
| 3 | 1000 | 1250 | 1500 | 250 |
| 4 | 1260 | 1575 | 1890 | 315 |
| 5 | 1588 | 1985 | 2382 | 397 |
| 6 | 2000 | 2500 | 3000 | 500 |
| 7 | 2518 | 2147.5 | 3777 | 629.5 |
| 8 | 3174 | 3967.5 | 4761 | 793.5 |
| 9 | 4000 | 5000 | 6000 | 1000 |
| 10 | 5000 | 6250 | 7500 | 1250 |

Note: f = frequency. f0 = fundamental frequency, i.e., the basis of the harmonic series of the triads' individual frequencies. Hz = Hertz.

**Supplementary Table 2.** Scheme for counterbalancing stimuli positions across sessions and participants.

| Participant | Session | Pair | Position 1 | Position 2 | Position 3 |
| --- | --- | --- | --- | --- | --- |
| *Bird A* | 1 | Ran vs VarReg | Silent | VarReg | Ran |
|  | 2 | Ran vs Reg | Reg | Ran | Silent |
|  | 3 | Reg vs VarReg | VarReg | Silent | Reg |
|  | 4 | Ran vs VarReg | Silent | Ran | VarReg |
|  | 5 | Ran vs Reg | Silent | Reg | Ran |
|  | 6 | Reg vs VarReg | Reg | Silent | VarReg |
|  | 7 | Ran vs VarReg | VarReg | Silent | Ran |
|  | 8 | Ran vs Reg | Reg | Ran | Silent |
|  | 9 | Reg vs VarReg | Silent | VarReg | Reg |
| *Bird B* | 1 | Ran vs Reg | Reg | Ran | Silent |
|  | 2 | Ran vs VarReg | Reg | Silent | VarReg |
|  | 3 | Reg vs VarReg | Silent | Reg | VarReg |
|  | [...] | [...] | [...] | [...] | [...] |

**Supplementary Figure 1.**
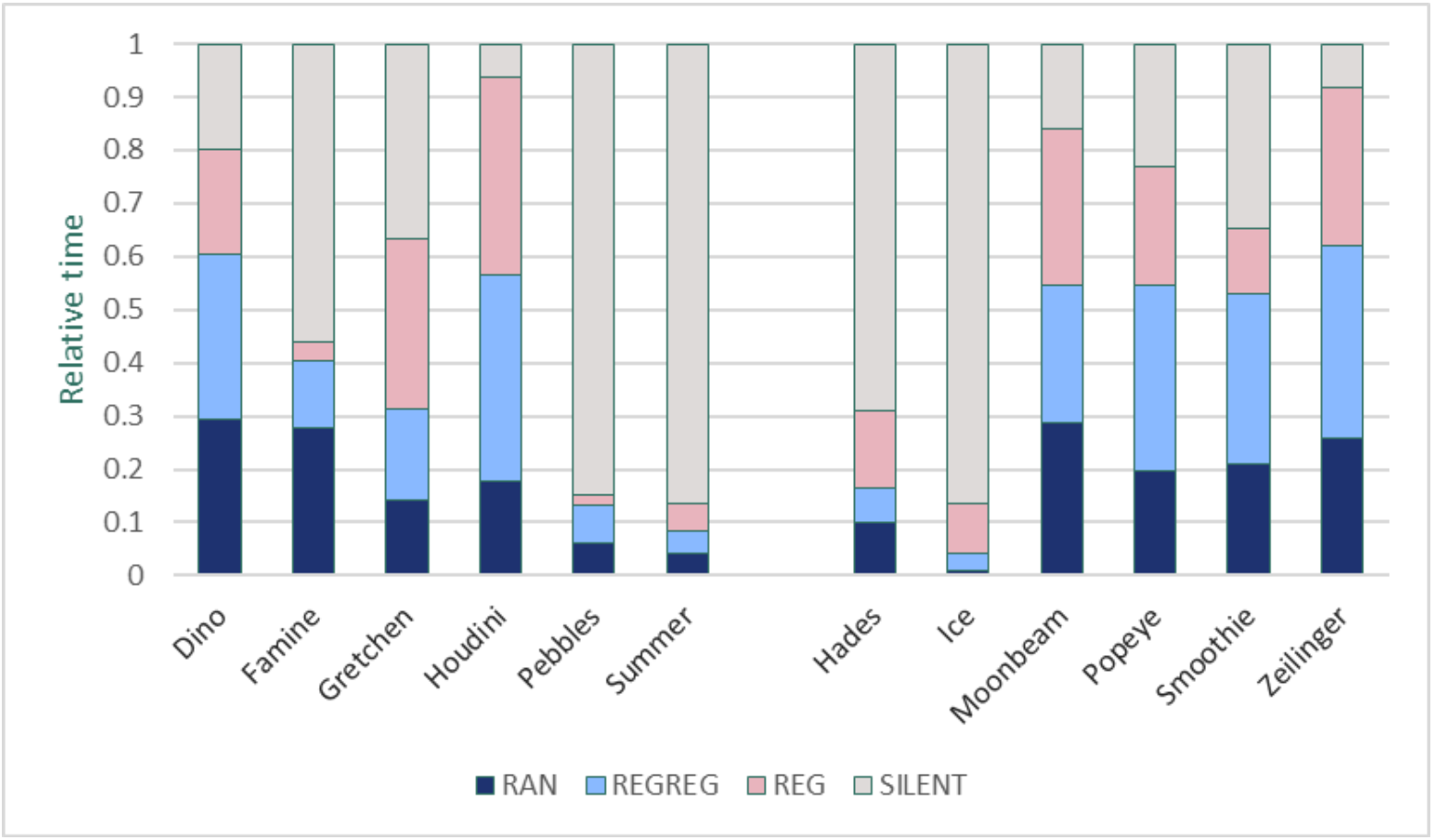
Individual relative mean average time the budgerigars spent with the different conditions. Note. Divided by the gap: Left females, right males.

**Supplementary Figure 2.**
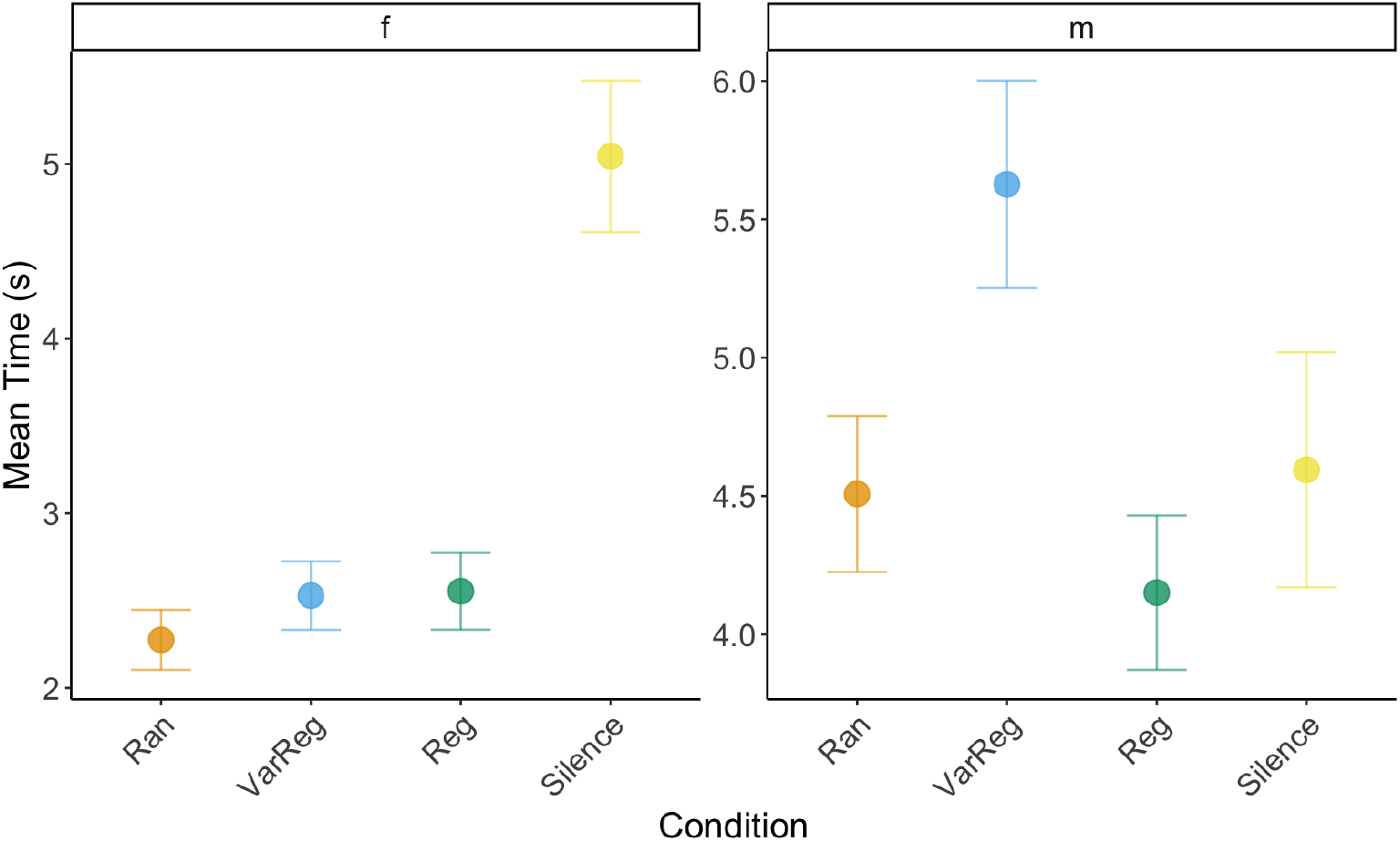
Mean time (in s) and standard error across human subjects and sessions for the conditions per trial in females (left) and males (right).

